# SSB facilitates fork substrate discrimination by PriA

**DOI:** 10.1101/2020.11.23.394411

**Authors:** Hui Yin Tan, Piero R. Bianco

## Abstract

PriA is a member of the SuperFamily 2 helicase family. Its role *in vivo* is to reload the primosome onto stalled replication forks resulting in the restart of the previously stalled DNA replication process. SSB is known to play key roles in mediating activities at replication forks and it is known to bind to PriA. To gain mechanistic insight into the PriA-SSB interaction, a coupled spectrophotometric assay was utilized to characterize the ATPase activity of PriA *in vitro* in the presence of fork substrates. The results demonstrate that SSB enhances the ability of PriA to discriminate between fork substrates 140-fold. This is due to a significant increase in the catalytic efficiency of the helicase induced by DNA-bound SSB. This interaction is species-specific as bacteriophage gene 32 protein cannot substitute for the *E.coli* protein. SSB, while enhancing the activity of PriA on its preferred fork, both decreases the affinity of the helicase for other forks and decreases catalytic efficiency. Central to the stimulation afforded by SSB is the unique ability of PriA to bind with high affinity to the 3’-OH placed at the end of the nascent leading strand at the fork. When both the 3’-OH and SSB are present, the maximum effect is observed. This ensures that PriA will only load onto the correct fork, in the right orientation, thereby ensuring that replication restart is directed to only the template lagging strand.

## Introduction

DNA replication is prone to various challenges that stall or delay the progression of forks (1). Challenges include damage to the template, a shortage of DNA synthesis precursors, secondary structure, and bound proteins (2–4). The repair of stalled replication forks frequently requires the actions of one or more DNA helicases (5). These critical enzymes harness the chemical, free energy of ATP hydrolysis to catalyze the unwinding of double-stranded DNA (dsDNA) (6,7). Many DNA helicases can act on unusual DNA structures such as Holliday junctions, stalled replication forks, and recombination intermediates (8–11).

Primosomal protein A (PriA) is one such DNA helicase that was originally identified as an essential factor required for the conversion of the complementary strand of ϕX174 to the replicative form during the initial stage of DNA replication (12,13). It is also required for bacteriophage Mu transposition and DnaA-independent replication of pBR322 (14,15). During the ϕX174 life cycle, PriA binds to a single-strand DNA (ssDNA) hairpin structure known as n’-primosome assembly site (PAS), leading to the subsequent assembly of the primosome, a complex responsible for primer RNA synthesis and duplex DNA unwinding at a replication fork (16,17). PAS sites also occur near the origin of pBR322 and can function as origins of DNA replication (18,19). In contrast, in the Mu life cycle, PriA directs the assembly of the preprimosome onto Mu forks following transpososome disassembly (15,20).

The 82kDa PriA protein consists of two domains (21,22). The N-terminal 181 aa are associated with DNA binding while the C-terminal 551 aa contains the ATP binding and DNA helicase motifs which are interrupted by two, C4-type zinc finger motifs (23). These Zn-finger motifs are essential for *in vitro* primosome assembly on PAS, for recombination-dependent DNA replication *in vivo*, and for interactions with other primosomal proteins (24–26). The DNA binding properties of PriA, mediated by the N-terminus, are consistent with its activity at stalled replication forks. It binds with high affinity to D-loops and to model, fork structures *in vitro* (20,27–29). This binding is mediated through specificity for DNA strands with accessible 3’-ends (27,30). Specificity is provided by a 3’-terminus binding pocket located in the OB-fold in the N-terminus of the protein (31).

PriA has been assigned to helicase SuperFamily 2 and has been shown to unwind DNA with a 3’→ 5’ polarity *in vitro* (32,33). DNA unwinding is fueled by the hydrolysis of ATP (dATP), is site-specific (*i.e.*, PAS), structure-specific, and ssDNA-dependent (34). Also, DNA unwinding of model fork substrates is stimulated by the single-stranded DNA binding protein (SSB) (35). This stimulation involves both a physical and functional interaction between the two proteins (36–39). As for several other proteins at the replication fork such as RecG, the physical interaction is mediated *via* the linker domain of SSB and the OB-fold in PriA (40).

Once bound to a stalled replication fork, PriA displays two types of activities. The 3’→5’ helicase activity is responsible for unwinding both the parental duplex ahead of the fork and the lagging-strand arm (20). The second activity is the loading of DnaB onto the lagging-strand template *via* a complex series of protein-protein interactions reminiscent of primosome assembly for ϕX174 DNA (17,20,41,42). The helicase activity of PriA is not required for this reaction (41). This leads to the loading of the replicative helicase, DnaB, from a DnaB-DnaC complex onto SSB-coated ssDNA. Once DnaB has been loaded, a new replisome forms, leading to the resumption of DNA replication (42,43).

It is becoming increasingly clear that SSB plays important roles at stalled replication forks in addition to binding of exposed ssDNA (44). It binds to the fork in a polar fashion and performs limited unwinding (45). SSB loads RecG and separately, PriA, and remodels these enzymes during the process (39,46). It ensures that Rep and UvrD do not process the same fork structure simultaneously (47). Finally, it stimulates the helicase activity of PriA (35,37). It was proposed that a combination of the fork structure and SSB would enhance the activity of PriA but the mechanism for this is unclear (35).

To understand the mechanism for these enhancements, a detailed characterization of the ATPase activity of PriA was performed in the presence of forks, and the catalytic efficiency of PriA in the presence and absence of SSB, determined. The data presented are consistent with previous work (20–22,27–31,35,48), and in addition, reveal that SSB increases the ability of PriA to discriminate the correct fork 140-fold relative to the incorrect forks. This is critical as there are a very small number of PriA molecules available in the cell and it is essential that the helicase does not mistakenly load on the incorrect strand or structure, and SSB ensures this will not happen. The outcome is that replication restart is directed to only the template lagging strand.

## Results

### PriA exhibits robust ATPase activity on ϕX174 ssDNA that is stimulated by SSB

The hydrolysis of ATP by PriA in the presence of various DNA molecules under several assay conditions was monitored utilizing a coupled spectrophotometric assay. We first analyzed the activity of the enzyme on ssDNA using M13 as the cofactor which is the standard for most of our helicase studies (47,49). The activity of PriA in the presence of this ssDNA was very low at 3.7±0.8 μM/min, consistent with previous work (Fig. 1A and (20,50)). Furthermore, SSB inhibits the ATPase activity of the protein 4-fold in the presence of this DNA cofactor. In contrast, SSB enhances the ATPase activity of RecG in the presence of M13 ssDNA (49).

**Figure 1.**
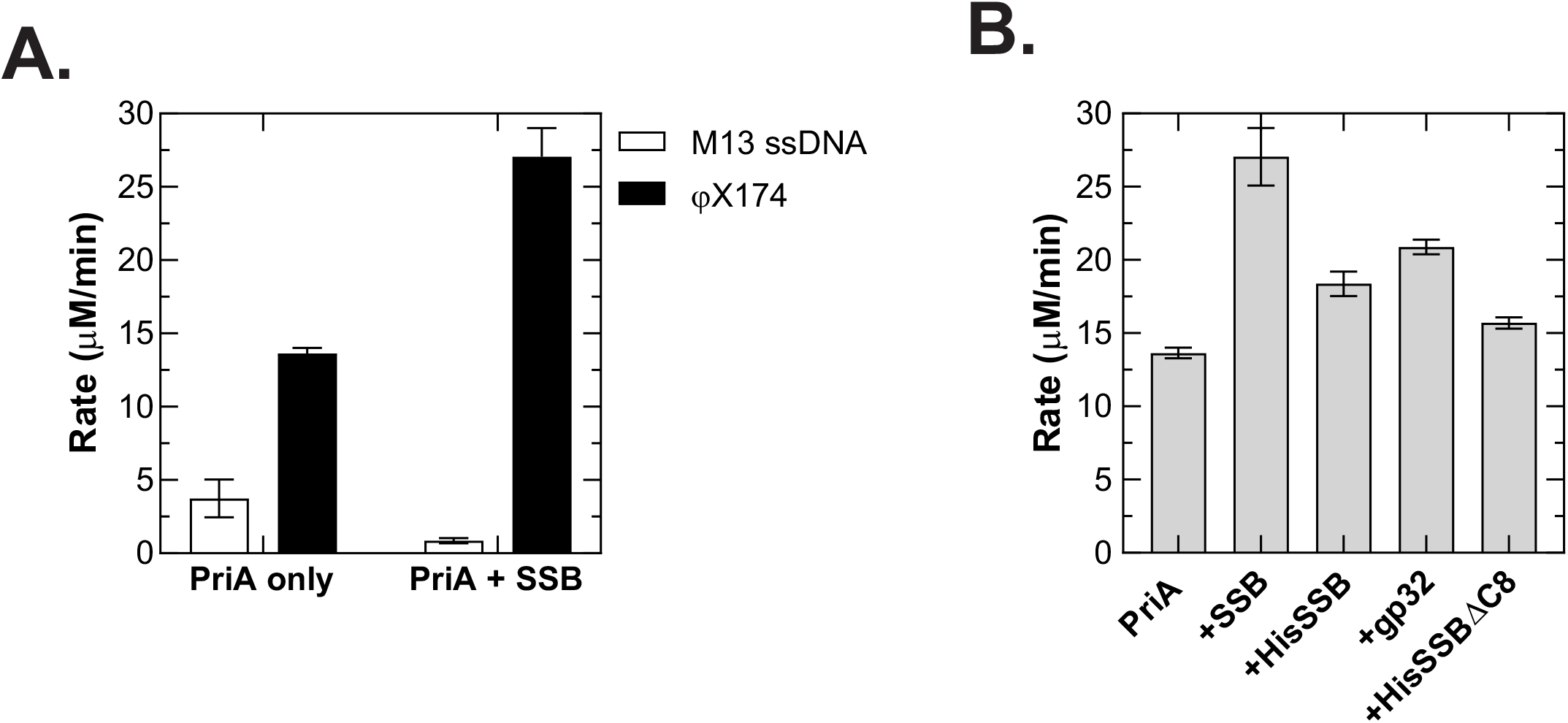
PriA exhibits ssDNA-dependent ATPase activity on ϕX174 DNA only. (A) SSB enhances the ATPase activity of PriA on ϕX174 but inhibits activity in the presence of M13 ssDNA. (B) Single strand binding proteins increase the ATPase activity of PriA in the presence of ϕX174 ssDNA. Assays were done as described in the Experimental Procedures and contained 10μM nucleotides of ssDNA, 20nM PriA, and 1μM single-strand binding proteins (where indicated). Reactions were initiated by the addition of PriA following a five-minute incubation of all other components at 37°C. Assays were done in duplicate on the same day.

PriA was initially identified as a factor that bound to a DNA hairpin structure in ϕX174 called PAS, for n’-<u>p</u>rimosome <u>a</u>ssembly <u>s</u>ite, leading to the subsequent assembly of the primosome, a complex responsible for primer RNA synthesis and duplex DNA unwinding at a replication fork (16,17). Therefore, we tested the ATPase activity of the helicase in the presence of ϕX174 ssDNA. The results show that the activity was 4-fold higher than that in the presence of M13 ssDNA (Fig 1A and (20,50)). Furthermore, the ATPase activity of PriA is enhanced 2-fold by the SSB protein consistent with previous work (50)). When compared to the SSB-containing reaction in the presence of M13 ssDNA, the ATPase activity of PriA is stimulated 32-fold. The higher level of activity of PriA in the presence of ϕX174 ssDNA is consistent with the helicase being a site-specific (*i.e.*, PAS), structure-specific, and ssDNA-dependent ATPase (34).

To determine whether the stimulation is specific to SSB, assays were repeated using different single-strand DNA binding proteins in the presence of ϕX174 ssDNA. The results show that the T4 gene 32 protein will substitute for SSB, although it is only 77% as effective (Fig. 1B). Second, the N-terminal histidine tag on SSB reduces the enhancement by 30%, but still stimulates the activity of the enzyme. Finally, there is a small but noticeable stimulation provided by SSBΔC8. As PriA and SSB interact physically and functionally, the result with SSBΔC8 indicates that in this assay, the mutant retains 85% of this interaction (20,37,38,51).

### SSB stabilizes PriA on ϕX174 ssDNA

Previous work has shown that SSB stabilizes the RecG DNA helicase on M13 ssDNA (49). This stabilization was observed as a two-fold increase in the salt-titration midpoint. To test if SSB has a similar effect on PriA, increasing [NaCl] was added to ongoing ATPase assays using ϕX174 ssDNA as the DNA cofactor. To permit a direct comparison to our published work, assays with RecG were repeated on the same day using the same assay components except that M13 ssDNA was the cofactor.

The results show that the salt-titration midpoint (STMP) for PriA alone was 41 mM and this increased 4-fold to 159 and 168mM in the presence of wild type and his-SSB, respectively (Fig 2A). SSBΔC8 produced a 1.7-fold increase in the STMP indicating that while this protein has defective C-termini, it stabilizes the helicase on ssDNA possibly through binding of the linker domain of SSB to the OB-fold in PriA (40). In contrast, the bacteriophage T4 gp32 protein does not affect the STMP. As expected, SSB also stabilizes RecG on ssDNA, producing a 2.8-fold increase in the STMP (Fig 2A). In contrast to PriA, SSBΔC8 had only a small but detectable effect on RecG. Therefore, SSB stabilizes both PriA and RecG on their respective ssDNA cofactors.

**Figure 2.**
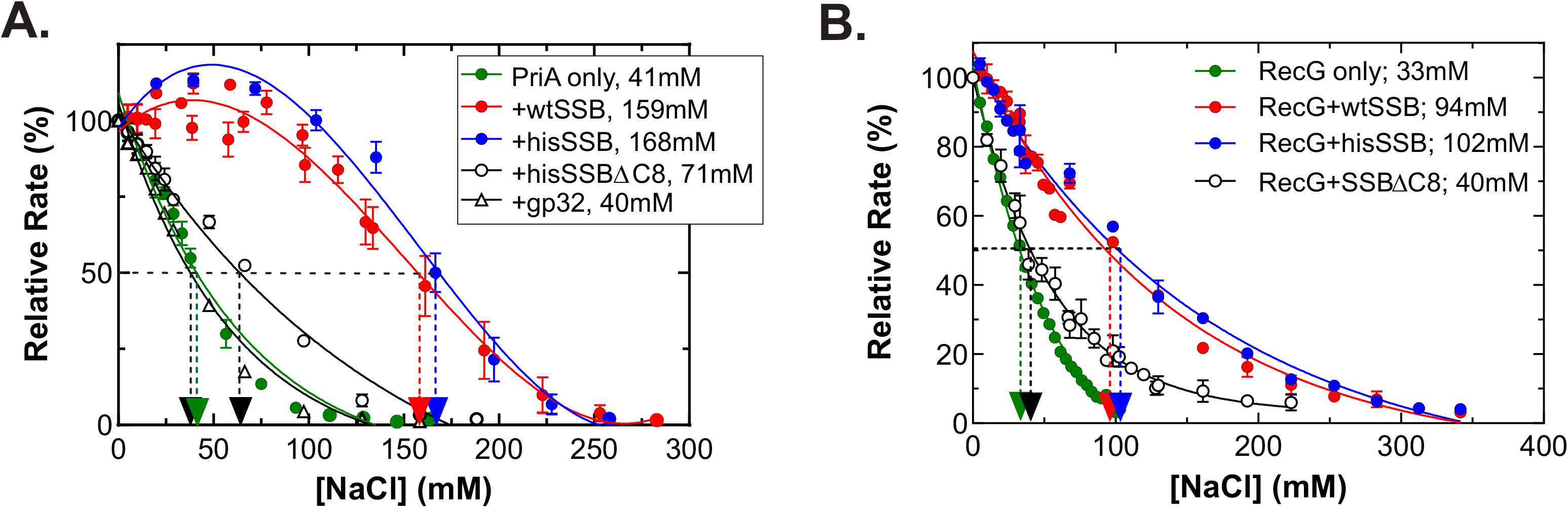
Only wild type E.coli SSB stabilizes fork rescue DNA helicases on ssDNA. (A) Stabilization of PriA on ϕX174 ssDNA and (B), Stabilization of RecG on M13 ssDNA. Assays were done as described in the Experimental Procedures and contained 10μM nucleotides of ssDNA, 20nM PriA or 10nM RecG, and 1μM single-strand binding proteins (where indicated). To obtain the salt-titration midpoint (STMP) the resulting rates of ATP hydrolysis at each concentration of NaCl were calculated during each phase of the reaction following the addition of NaCl, and expressed as a percent of the reaction rate in the absence of added NaCl. The dashed lines indicate the STMP for each reaction. A minimum of four separate assays were done for each reaction condition. The STMP data for RecG have been published earlier, but assays were redone and the resulting data presented here for direct comparison to PriA (49,70).

### SSB affects the ATPase activity of PriA in a fork structure-dependent manner

Previous work from several laboratories showed that SSB interacts with PriA both physically and functionally (20,22,37–39,51). To understand the mechanism of these interactions at forks where PriA plays critical roles *in vivo*, we utilized a series of model fork substrates to characterize the ATPase activity of PriA. We previously used these forks to characterize the ATPase activity of other DNA helicases - RecG, RuvAB, Rep, and UvrD (47,49,52).

These model forks, shown schematically in Table I, are formed by annealing purified oligonucleotides to produce a fork with flayed ends (fork 1); a fork with a gap in the lagging strand arm (fork 2 which also has a 3’-OH positioned at the fork on the leading strand arm); fork 3 (which has a gap in the leading strand); a fork with two duplex arms (fork 4) and finally a Holliday Junction (fork 5). Forks 1-3 which contain one or more ssDNA arms comprise group I and are thought to mimic nascent, stalled replication fork structures. Fork 4 and the HJ, which contain duplex DNA arms, are assigned to group II as they are thought to mimic regressed fork structures. These last two forks were included initially to permit a comparison to RecG and RuvAB, even though PriA does bind to them but will not unwind them unless they contain a 5nt gap at the fork (20,27,35,49,52). At the center of each fork is a homologous core of 24bp flanked by heterologous sequences so that PriA can mediate the unwinding of each of the substrates (data not shown and (53–55)).

**Table I.**
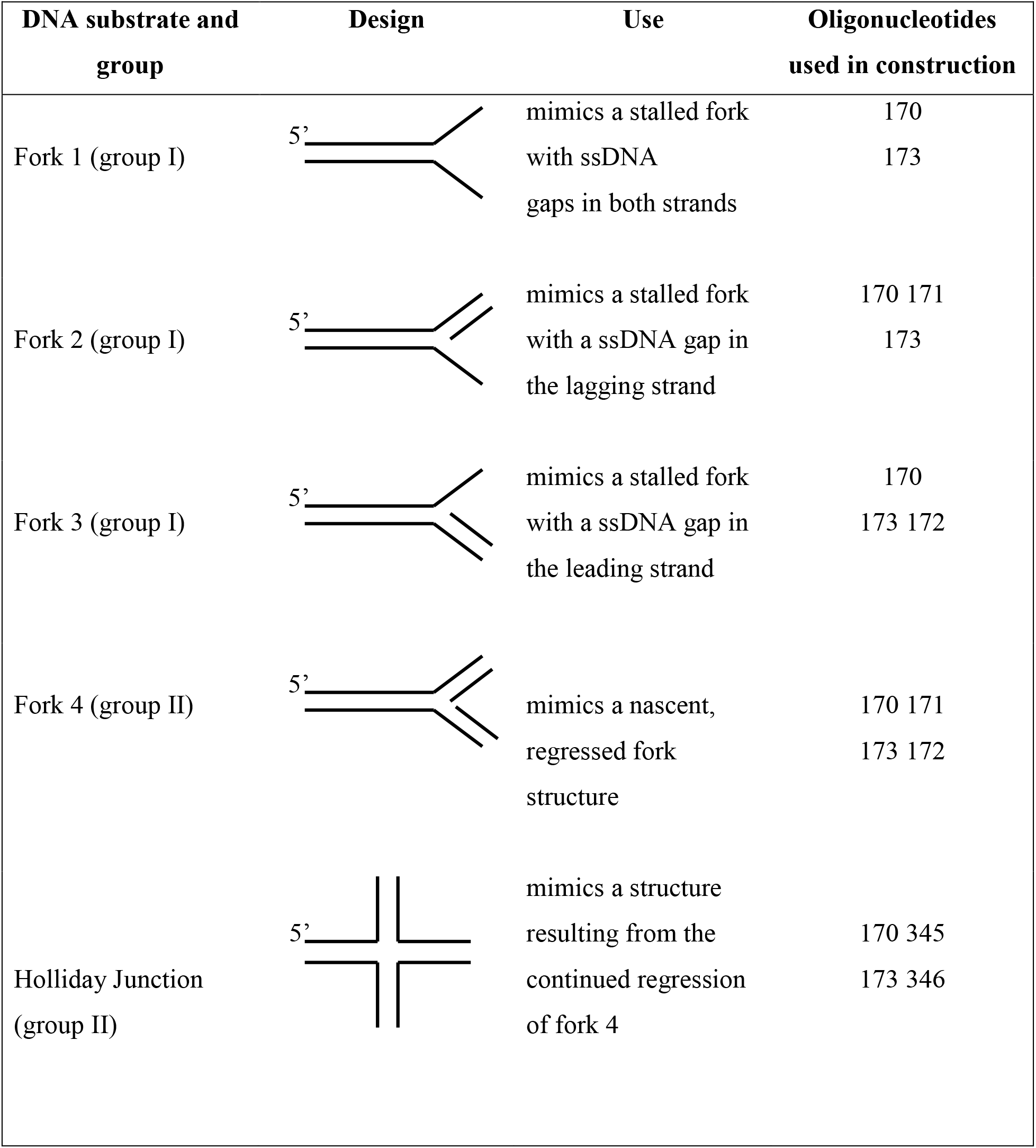
DNA cofactors and their uses.

| DNA substrate and group | Design | Use | Oligonucleotides used in construction |
| --- | --- | --- | --- |
| Fork 1 (group I) |  | mimics a stalled fork with ssDNA gaps in both strands | 170<br>173 |
| Fork 2 (group I) |  | mimics a stalled fork with a ssDNA gap in the lagging strand | 170 171<br>173 |
| Fork 3 (group I) |  | mimics a stalled fork with a ssDNA gap in the leading strand | 170<br>173 172 |
| Fork 4 (group II) |  | mimics a nascent, regressed fork structure | 170 171<br>173 172 |
| Holliday Junction (group II) |  | mimics a structure resulting from the continued regression of fork 4 | 170 345<br>173 346 |

Previous studies of DNA helicase activity were done using model Holliday junctions and stalled fork substrates which themselves are influenced by magnesium ion concentration and could affect the resulting activity of PriA accordingly (47,49,56–60). Therefore, to understand the mechanism of PriA interactions at forks, we first assessed the ATPase activity of the helicase as a function of magnesium ion concentration (Fig. 3A). The data show that the activity of PriA was maximal between 1 and 5mM, with the highest level of activity being observed in the presence of fork 2, which mimics a fork with a gap in the nascent lagging strand and has a critical 3’-OH group at the fork (30). This is followed by fork 1 which has two single-stranded arms and the Holliday junction (HJ). Extremely low levels of activity were observed in the presence of fork 3, which mimics a fork with a gap in the nascent leading strand. The ATPase activity was 13-fold lower at the optimal [MgOAc]. The fork preference exhibited here is consistent with previous work showing that the preferred fork for PriA has a gap in the nascent lagging strand. However, it was surprising to us that PriA exhibited high levels of ATPase activity in the presence of fork 4 (a fork with two duplex arms) and the Holiday junction. This follows because it was shown that while PriA can bind these DNA structures, their unwinding of these DNAs required the presence of a 5nt gap in the fork arms (30,35). Fork 4 and the HJ were not studied further.

**Figure 3.**
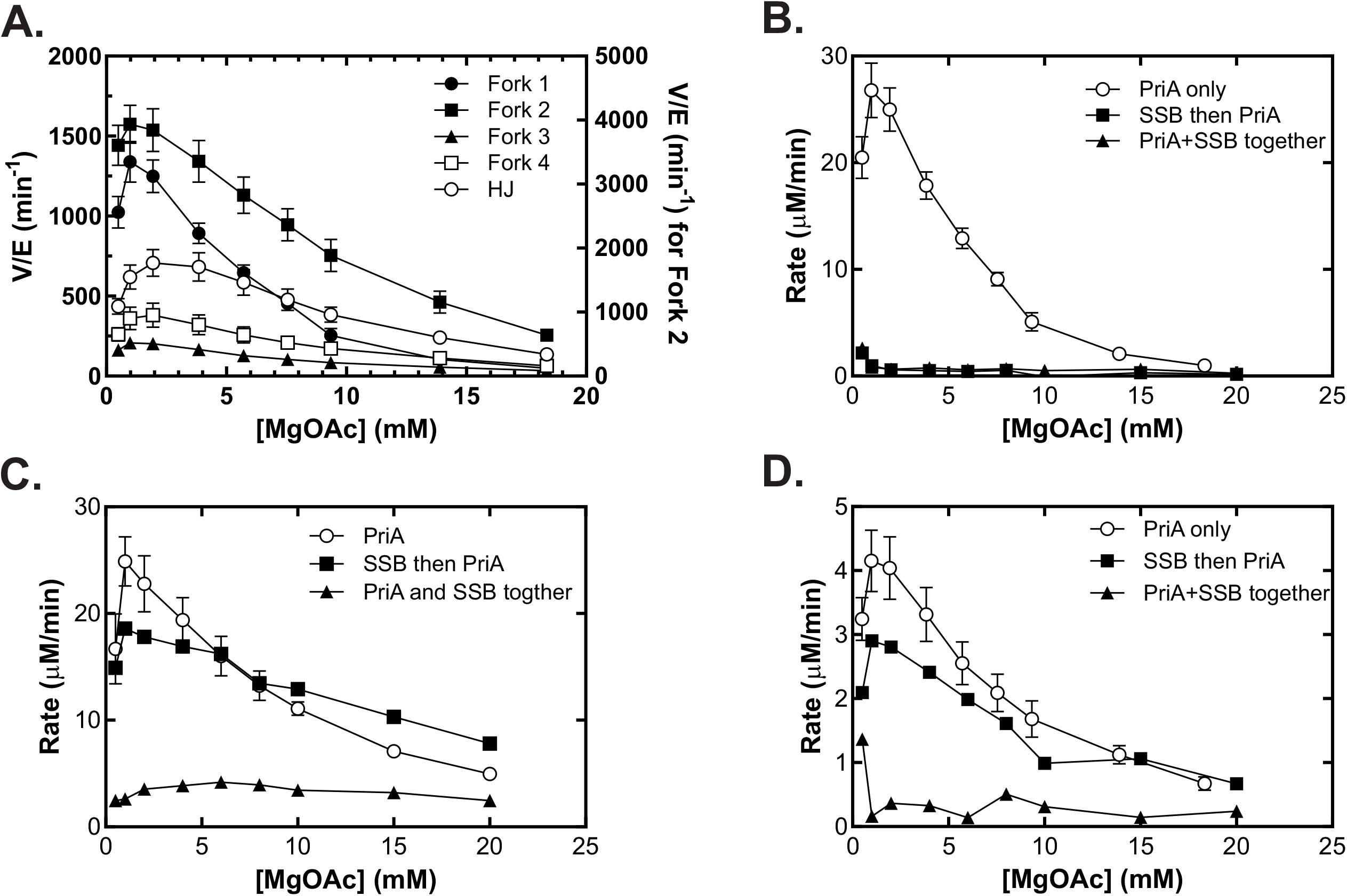
The order of addition dictates the effects of SSB on PriA in the presence of stalled fork DNA cofactors. Magnesium acetate titrations were done using fork cofactors as indicated. Assays contained 10 nM PriA helicase, 1 mM ATP, and 100nM molecules of each DNA cofactor. (A) The magnesium optimum for PriA is fork-structure dependent. (B) SSB inhibits the ATPase activity of PriA in the presence of a fork with two single-stranded tails. (C and D), when added first, SSB does not inhibit PriA in the presence of forks with a gap in the nascent lagging (C) or leading strands(D).

To understand how SSB influences the ATPase activity of PriA in the presence of forks with single-strand character, MgOAc titrations were repeated but in the presence of SSB and the data compared to that observed for PriA alone (Fig. 3B-D). For fork 1, which has 2 ssDNA arms, the presence of SSB virtually eliminated the ATPase activity of PriA (Fig 3B). The inhibition seen here is greater than the 4-fold effect seen previously (35). Further, inhibition was independent of whether SSB (200nM tetramer) was added to the forks first or allowed to bind to PriA (10nM) before being added to the reaction. Also, inhibition was specific to *E.coli* SSB with wild type having the greatest effect on PriA (Supplementary Figure 1A). Even SSBΔC8 which has mutant C-termini, is effective at inhibiting the ATPase activity of PriA on fork 1. In contrast, bacteriophage T4 gp32, which binds to ssDNA with a polarity opposite to that of SSB and is not known to bind PriA, stimulates the ATPase activity of PriA on fork 1 (45).

In contrast to fork 1, SSB (100nM tetramer) when added to forks 2 or 3 before PriA, had only a minimal effect on the ATPase activity of the helicase at the magnesium optimum (Fig. 3C and D). Surprisingly, when PriA and SSB were premixed, ATPase activity in the presence of forks 2 and 3 was inhibited several-fold, independent of the concentration of MgOAc. This was specific to wild type SSB for fork 2 and occurred with all single-strand binding proteins tested for fork 3 (Supplementary Figure 1). In summary, the data in this section show that when added separately, SSB stimulates the ATPase activity of PriA on fork 2 the preferred DNA cofactor, while it inhibits the ATPase activity of PriA on forks 1 and 3, which are not the preferred forks.

### ATP titrations reveal insight into how SSB regulates PriA at forks

To further understand how single-strand binding proteins (ssbs) influence the activity of a DNA helicase at a fork, we performed ATP titrations and determined the relevant kinetic parameters for PriA. Assays were done in the presence of forks 1-3 and were done with PriA only, and separately in the presence of either SSB, SSBΔC8 or gp32. The raw data are shown in Supplementary Figure 2, kinetic parameters are presented in Table II and the final analysis is shown in Figure 4.

**Table II.** ATP kinetic parameters for PriAa.

| Proteins present | DNA | $K_m^{ATP}$<br>(μM) | $V_{max}$<br>(μM/min) | $k_{cat}$<br>($\text{min}^{-1}$) | $k_{cat}/K_m$<br>($\text{min}^{-1}/\text{nM}$) |
| --- | --- | --- | --- | --- | --- |
| PriA | Fork 1 | 2,740±936 | 8.1± 1.3 | 810 | 0.3 |
| | Fork 2 | 882±232 | 16.9±1.5 | 1,690 | 1.9 |
| | Fork 3 | 1,123±635 | 2.2±0.3 | 220 | 0.2 |
| PriA+SSB | Fork 1 | 1,303±551 | 43± 5.6 | 4,300 | 3.3 |
| | Fork 2 | 852±158 | 144±10 | 14,400 | 16.9 |
| | Fork 3 | 1,814±884 | 5.1±0.7 | 510 | 0.3 |
| PriA+SSBΔC8 | Fork 1 | 1,871±664 | 81± 13 | 8,100 | 4.3 |
| | Fork 2 | 1,733±729 | 67±13 | 6,700 | 3.9 |
| | Fork 3 | 4,540±1,077 | 31±4 | 3,100 | 0.7 |
| PriA + T4 gp32 | Fork 1 | 4,065±1,256 | 144± 21 | 14,400 | 3.5 |
| | Fork 2 | 6,479±2,283 | 110±22 | 11,000 | 1.7 |
| | Fork 3 | 2,677±2051 | 15±2 | 1,500 | 0.6 |

a. Assays were done as described in the Materials and Methods. The data were approximated by the Michaelis-Menten equation.

**Figure 4.**
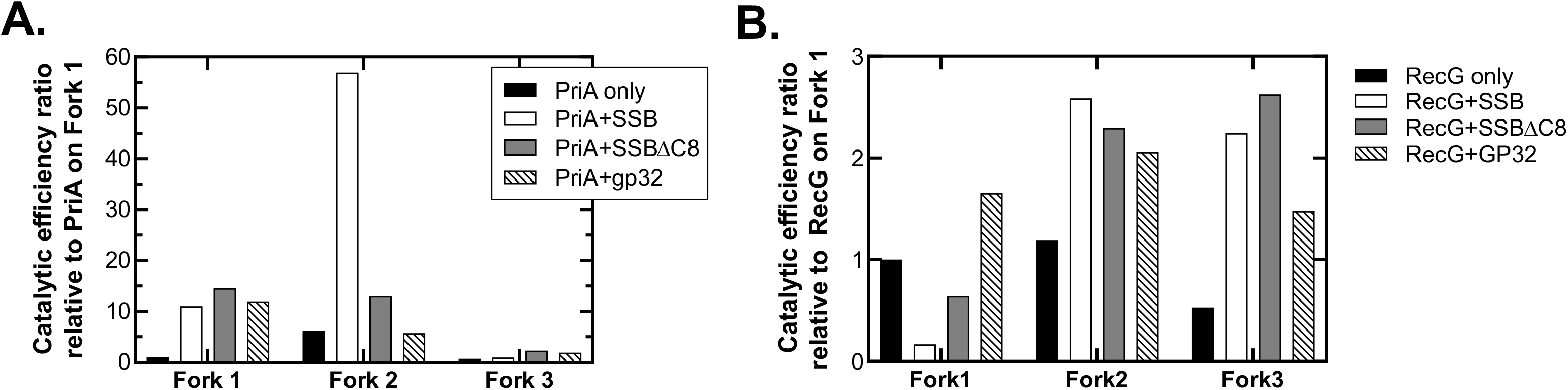
Wild type *E.coli* SSB enhances fork substrate-discrimination by PriA. To enable direct comparison, the data in each panel have been normalized to the catalytic efficiency for the ATPase activity of each helicase alone in the presence of fork 1. Kinetic data were obtained from ATPase assays and are shown in graph format in Supplementary Figure 2 for PriA (not shown for RecG). The values are presented in Table II and were used to calculate the ratios presented. (A), Wild type SSB specifically increases the catalytic efficiency of PriA in the presence of fork 2, its preferred fork cofactor. (B), SSB has minimal effects on the catalytic efficiency of RecG. Assays with each fork were done as ATP titrations in the absence and presence of single-strand binding proteins.

First, an inspection of the curves in Supplementary Figure 2A shows that under these assay conditions, the preferred fork substrate is fork 2 which has a gap in the nascent lagging strand and the requisite 3’-OH group in the nascent leading strand, consistent with previous work (30,35,61). Second, SSB increases the V_max_ of PriA in the presence of forks one to three, 5-, 9- and 2.3-fold, respectively (Supplementary Figure 2B and for precise numbers see Table II). Third, SSBΔC8 also enhances the ATPase activity of PriA. However, it enhances the V_max_ of PriA 10-fold on fork 1, 14-fold on fork 3, and only 4-fold on fork 2 (Supplementary Figure 2C and Table II). Fourth, gp32 also stimulates the V_max_ of PriA in the presence of forks, but the stimulation is highest in the presence of fork 1 (18-fold), 6.8-fold for fork 3, and only 6-fold for fork 2 (Supplementary Figure 2D and Table II). In summary, gp32 enhances V_max_ highest on fork 1 whereas SSBΔC8 affects forks 1 and 3 equally well. They do not show a fork 2-specific effect. In contrast, this initial analysis suggests that only SSB significantly enhances the activity of PriA in the presence of its preferred fork substrate (fork 2).

To further understand the mechanism for this SSB- and fork-specific stimulation, kinetic parameters were calculated and the catalytic efficiency of the helicase assessed (Table II). First, the catalytic efficiency of PriA alone in the presence of fork 2 is 1.9, and this is 10-fold higher than fork 3 and 6-fold higher than fork 1 (Table II). This is consistent with the requirement for a 3’-OH group at the fork on the nascent leading strand, in the full activation of the ATPase activity of the enzyme (30). The high catalytic efficiency observed for this fork DNA is a combination of elevated V_max_ and a low K_m_^ATP^. In the absence of the 3’-OH group at the fork, V_max_ decreases 2-to 8-fold and this is accompanied by an increase in the K_m_^ATP^ as seen for forks 1 and 3.

In the presence of SSB, the catalytic efficiency of PriA changes dramatically. First, it increases 9-fold on fork 2, 10-fold on fork 1, and is unchanged on fork 3, relative to PriA alone on these same forks (Table II). SSBΔC8 increases the catalytic efficiency 14-fold on fork 1, 2-fold on fork 2, and 3.5-fold on fork 3. T4 gp32 also increases catalytic efficiency 12-fold on fork 1, 3-fold on fork 3 but does not affect the catalytic efficiency of PriA in the presence of fork 2. In summary, all 3 ssbs stimulate catalytic efficiency equally well in the presence of fork 1 (10-to 14-fold), and also for fork3, although the effect is much less (2-to 4-fold). In contrast, the stimulation in catalytic efficiency in the presence of fork 2 observed for SSB is 4-fold higher than SSBΔC8 and 10-fold higher than gp32.

As the presence of the 3’-OH group increases the catalytic efficiency of PriA 6-to 10-fold in the presence of fork 2 relative to forks 1 and 3 respectively, the effects of SSB on catalytic efficiency in the presence of fork 2 were recalculated relative to that of PriA alone on fork 1 to ascertain the combined effect. When analyzed in this manner, the effect of the 3’-OH group and SSB increases the catalytic efficiency of PriA 56-fold in the presence of fork 2 (Fig. 4A). When calculated in the same way, SSBΔC8 has a 13-fold enhancement and gp32 is 5.7-fold. However, these two proteins increase the catalytic efficiency of PriA on fork 1, 14.5- and 12-fold respectively. Thus, while SSBΔC8 does affect k_cat_/K_m_^ATP^, this effect is mediated through its ability to bind ssDNA. In contrast, when the 3’-OH is present, gp32 inhibits the catalytic efficiency of the helicase indicating then when an ssb binds in ssDNA in the opposite orientation, PriA ATPase is impaired. Furthermore, when the 3’-OH group is absent as it is in fork 1, any single-strand binding protein will increase the catalytic efficiency of PriA (Fig. 4A). This enhancement is due to the ssb binding to the lagging strand arm. This follows because when the lagging strand arm is duplex, single-strand binding proteins have only a minimal effect as observed for fork 3 (Table II and Fig. 4A).

Collectively, these data show that there are two components to the catalytic efficiency enhancement induced by SSB. First, there is direct ssDNA binding and second, there is the combination of the ssb-PriA interaction and the 3’-OH group, with only SSB providing the maximal stimulation for the latter. Interactions between the SSB linker and the PriA OB-fold are likely involved in this stimulation (40). The large stimulation afforded by SSB on fork 2 is a result of an 8.5-fold increase in V_max_ (Table II). In contrast, SSB does not increase the catalytic efficiency of PriA in the presence of fork 3. Even though V_max_ increases (Supplementary Figure 2B and Table II), this is accompanied by a corresponding increase in the K_m_^ATP^, resulting in an unchanged value for k_cat_/K_m_^ATP^. Consequently, the enhancement in catalytic efficiency observed for PriA in the presence of fork 2 and SSB, relative to PriA alone in the presence of fork 3 is 84.5-fold.

In contrast to PriA, the effects of single-strand binding proteins on the catalytic efficiency of RecG are at best, modest (Fig. 4B). SSB does however decrease the catalytic efficiency of RecG in the presence of fork 1 while increasing this kinetic parameter two-fold for fork 2 and five-fold for fork 3 (the preferred fork substrate for this enzyme; (49,52)). However, comparable effects are also observed for SSBΔC8 and gp32 indicating that the presence of a single-strand binding protein enhances the catalytic efficiency of RecG but does not facilitate further substrate discrimination. These data are consistent with a special and unique interaction between SSB and PriA. In summary, these kinetic data show that while PriA has a known substrate preference for a fork with a 3’-OH group at the fork and a gap in the nascent lagging strand, SSB, bound to this single strand arm, significantly enhances fork substrate discrimination by PriA.

### Analysis of DNA kinetic parameters confirms the DNA cofactor specificity

To determine whether SSB influences the binding specificity of PriA for fork substrates, ATPase assays were repeated but this time the concentration of DNA was varied and kinetic parameters calculated. The results show that as anticipated, the preferred cofactor is fork 2 as the catalytic efficiency of the enzyme is highest in the presence of this DNA (Table III). It was 1.7-fold higher than for fork 1 and 30-fold higher than that observed for fork 3. We note for fork 3, cofactor inhibition was observed for this fork (Supplementary Figure 3). Furthermore, for fork 2, a Hill coefficient of 2.1±0.3 was observed suggesting that under these conditions, PriA can bind at least two of these forks resulting in high levels of activity. The values for K_m_^app, DNA^ obtained here are comparable to the K_d_ values reportedly previously for comparable fork substrates (27,30).

**Table III.** Kinetic DNA parameters for PriAa.

| DNA | $K_m^{app}$<br>(nM) | $V_{max}$<br>(μM/min) | $k_{cat}$<br>(min<sup>-1</sup>) | $k_{cat}/K_m^{app}$<br>(min<sup>-1</sup>/nM) | Hill<br>Coefficient |
| --- | --- | --- | --- | --- | --- |
| Fork 1 | 5.7±0.7 | 3.6± 0.1 | 1,800 | 316 | 1.2±0.1 |
| Fork1 +SSB | 13.4±3.3 | 3.3±0.2 | 1,650 | 124 | 1.0 |
| Fork 2 | 6.8±0.5 | 7.3±0.2 | 3,650 | 537 | 2.1±0.3 |
| Fork2+SSB | 5.3±0.7 | 4.5±0.1 | 2,250 | 424 | 1.0 |
| Fork 3<sup>b</sup> | 17±11 | 0.6±0.04 | 300 | 18 | 1.9±0.5 |
| Fork3+SSB | ND | ND |  |  |  |

a. Assays were done as described in the Experimental Procedures. The data were approximated by the Hill equation.
c. ND, not done.

When assays were done in the presence of stoichiometric SSB relative to forks, several changes in the kinetic parameters of PriA were observed. First, the Hill coefficient for DNA binding for each fork was 1. Second, the catalytic efficiency of the enzyme in the presence of fork 1 decreased 2.5-fold. This is attributed to a two-fold decrease in the apparent affinity of the enzyme for this fork from 5.7±0.7 to 13.4±3.3 nM (Table II). Thus, this is the third contribution to further enhancing substrate discrimination by PriA – lowering the apparent affinity for a DNA cofactor which is not ideal for replication restart. For fork 2, there was a 1.2-fold decrease in k_cat_/K_m_^app^ and this was due to the decrease in V_max_. However, SSB still produced a 2-fold increase in the ability of PriA to discriminate between forks 2 and 1, due to the large difference in catalytic efficiency. Therefore, the combined effects of contributions of SSB on the ability of PriA to discriminate between forks is 140-fold.

## Discussion

The primary conclusion of this study is that SSB facilitates fork discrimination by PriA by as much as 140-fold. SSB exerts these effects on both the ATP and DNA components of the helicase, with significant increases in catalytic efficiency being observed relative to those determined in its absence. The effects of SSB are greatest when PriA is bound to its preferred fork which has a gap in the lagging strand and a 3’-OH group positioned at the fork on the nascent leading strand. The combination of these effects, as well as the blocking of PriA binding to aberrant single-strands of DNA exposed at forks, ensures that the preprimosome is loaded onto the template lagging strand and that replication restart proceeds in the correct direction.

PriA is an unusual DNA helicase with unique DNA binding specificity that is in some respects similar to RecG but very distinct in its own right (23,49,52,62). RecG binds to D-loops and prefers a fork with a gap in the nascent leading strand. PriA also binds to D-loops and model fork structures but demonstrates a preference for a fork with a gap in the nascent lagging strand (20–22,27–29,35,48). It was later shown that the enzyme has a 3’-terminus binding pocket that plays a key role in facilitating specific binding to a fork when this 3’-OH is positioned on the nascent leading strand at the fork producing a K_d_ = 1-2 nM (30,31). This binding is also critical to the activation of the ATPase activity (29). When this 3’-OH group is absent or, blocked by the addition of a phosphate group, K_d_ increases 8-to 10-fold and, while ATPase activity is still observed, it is induced to a 5-fold lower level.

We have extended these studies and show that indeed the 3’-OH group positioned at the fork on the nascent leading strand, is essential for efficient ATPase activity of PriA (Figs. 3 and 4). This group enhances the catalytic efficiency of the enzyme 6-fold (fork 2) relative to forks where it is absent (forks 1 and 3). This results from a 3-fold lower K_m_^ATP^ and a 2-fold higher V_max_ in DNA titrations (Tables II and III). Thus, this small group, when correctly positioned to bind to the 3’-terminus binding pocket of PriA, has a dramatic effect on the ability of the enzyme to hydrolyze ATP. But that is not all.

In this study, the dramatic effects of SSB on the activity of PriA are demonstrated. The total of these effects is a minimum, 140-fold increase in catalytic efficiency of PriA on its preferred fork substrate relative to PriA alone in the presence of other forks. The stimulation is provided only by SSB and to a lesser extent, SSBΔC8 but not by T4 gp32 (Fig. 4). This follows because there are two SSB components to the catalytic efficiency enhancement. First, there is direct ssDNA binding to the lagging strand template and second, there is an interaction between the SSB and PriA that involves linker/OB-fold binding (40,45). Binding of SSB to the lagging strand blocks access to this ssDNA thereby ensuring that PriA can only bind to the fork in one orientation. This makes sense because PriA binds to ssDNA with high affinity (27,29). When the 3’-OH group present, high-affinity binding occurs (30,31). Second, SSB binds directly to PriA and loads the enzyme onto the DNA (37–39). In the process, SSB impacts both the DNA binding and ATPase kinetics of PriA. It is unknown whether the SSB-PriA or, SSB-ssDNA binding transmitted to the helicase or, a combination of both interactions, are responsible for the dramatic enhancement in catalytic efficiency. This requires the 3’-OH group bound at the fork and SSB, consistent with the proposal by Jones and Nakai (35). The outcome of the presence of a fully functional SSB bound to the fork is a dramatic increase in the ability of PriA to discriminate between fork structures so that it binds to, and processes, only the correct structure.

SSB, in addition to enhancing substrate discrimination by PriA, also stabilizes the enzyme on the DNA as evidenced by the 4-fold increase in the STMP (Fig. 2A). This indicates that once loaded and translocation and unwinding ensue, PriA will remain bound to the template lagging strand. SSB binds to ssDNA with high affinity and 10pN of force are required to displace a single tetramer (63,64). But, it does not represent an impassable block to the translocating helicase as shown previously (36). Thus, like RecG, SSB loads PriA onto the DNA and can be subsequently displaced during translocation by the helicase (39,46,65,66).

In exponentially growing cells, there are more than 2,000 SSB tetramers per cell (67). In these same cells, there are on average 2-4 DNA replication forks per cell with as many as 25 tetramers bound per fork. At each fork, there is 0.5 to 1kb of ssDNA available (68). Using a site size of 40 nucleotides occluded per tetramer, there would be on average 25 tetramers bound per fork or 100 per cell with the free SSB localizing to the inner membrane (69). In contrast to SSB, the levels of PriA are significantly lower at 2-4 molecules per cell (67). By binding to SSB, PriA also localizes to the inner membrane in the absence of exogenous DNA damage (38). Thus, it is not unreasonable to propose that when forks stall, PriA must be transferred to the DNA. However, results herein show that premixing SSB and PriA prior to adding these to the DNA, reduces the activity of PriA at forks (Fig. 3B-D). If PriA-SSB complexes did form before being added to reactions then this form of the helicase is either inactive or it was a titration effect producing the observed inhibition. If the former is correct, it suggests that PriA must be transferred from the storage form complex, to the SSB complexes already bound at the fork. The mechanism for this is unknown but could involve SSB to SSB transfer. Once there, SSB plays its important roles in effecting the outcome of events at a fork as shown previously for Rep, UvrD and for RecG (40,46,47,52,70). This makes sense because SSB loads PriA onto the DNA and remodels the helicase in the process and there is also a functional interaction between SSB and PriA (20,35–40,51). For PriA a significant component of this set of interactions is the 140-fold increase in the ability of the enzyme to discriminate the correct fork substrate from the incorrect one. Due to the very small number of PriA molecules available in the cell it is essential that a mistake not be made and SSB ensures this will not happen.

The results herein have elucidated the effects of the physical and functional interactions of SSB on PriA and these are rather complex. Part of this complexity arises because ssbs bind ssDNA in a polar fashion and can partially unwind duplex DNA at the fork (45). Second, single-stranded DNA binding by SSB plays a role in directing PriA to bind the fork in the correct orientation (35). Third, binding of the ssb to PriA is important as the effects observed are restricted to *E.coli* SSB and to a lesser extent SSBΔC8. This is likely afforded by binding of the linker domain of SSB to the OB-fold present in the N-terminus of PriA (40). Fourth, the structure of the DNA on which activity is being observed, that is to which PriA must bind, is also a key factor as suggested previously and shown herein (20,27,35,50). This follows because the maximal stimulation in both ATP- and DNA titrations by SSB was observed for fork 2. On this fork, pre-bound SSB directs PriA to its correct position where it binds with high affinity to the fork taking advantage of the 3’-terminus binding pocket docking with the 3’-OH positioned at the end of the nascent leading strand at the fork. This ensures that the preprimosome is loaded onto the correct strand at the fork (that is the template lagging strand) so that the resumption of DNA replication proceeds in the right direction.

### Experimental Procedures

### Materials

All chemicals were reagent grade, made up in Nanopure water and passed through 0.2_μm pore size filters. Yeast Extract and tryptone were from Becton Dickinson and Company (MD, USA). NaCl, sucrose, tris-base, KCl, Na_2_HPO_4_, NaH_2_PO_4_, EDTA, acetic acid, methanol and nickel sulfate were from J.T. Baker (NJ, USA). Ampicillin was from Fisher (NJ, USA). IPTG was from OmniPur (NJ, USA). Kanamycin, chloramphenicol, lysozyme and sodium deoxycholate were from Sigma (MO, USA). Benzonase was from Novagen (NJ, USA). Imidazole was from EMD (NJ, USA). Coomassie brilliant blue R-250 was from Bio-Rad Laboratories (CA, USA). Glucose was from Mallinckrodt (KY, USA). Nonidet P40 substitute was from USB (OH, USA). ATP, DEAE Sepharose Fast Flow, Q-Sepharose, the HisTrap FF, the 16/10 heparin FF, Mono Q and the Mono S 5/50 GL columns were from GE Healthcare Life Sciences (NJ, USA). Phosphoenol pyruvate (PEP), nicotinamide adenine dinucleotide (NADH), pyruvate kinase (PK), lactate dehydrogenase (LDH) and the ssDNA-cellulose resin were from Sigma. Phosphocellulose (P11) was from Whatman. Bio-Gel^®^ HTP hydroxylapatite was from Bio-Rad. Dithiothreitol (DTT) was from Acros Organics. BSA and *Hind*III were purchased from New England Biolabs. Wheat Germ Topoisomerase I (WGT) was from Promega.

### Reagents

All solutions were prepared using Barnstead Nanopure water. Stock solutions of Phosphoenol pyruvate (PEP) were prepared in 0.5 M Tris-acetate (pH 7.5). ATP was dissolved as a concentrated stock in 0.5 M Tris-HCl (pH 7.5), with the concentration determined spectrophotometrically using an extinction coefficient of 1.54 × 10^5^ M^−1^ cm^−1^. NADH was dissolved in 10 mM Tris-acetate (pH 7.5), concentration determined using an extinction coefficient of 6,250 M^−1^ cm^−1^, and stored in small aliquots at −80°C. Dithiothreitol (DTT) was dissolved as a 1M stock in nanopure water and stored at −80°C. All reaction buffers described below were assembled at 10 times reaction concentration and stored in 1mL aliquots at −80 °C.

### DNA cofactors

For all DNA cofactors, the concentrations of stock solutions were determined in μM nucleotides using the extinction coefficients as indicated below. To permit direct comparisons between fork DNA cofactors, concentrations and subsequent K_m_^DNA,app^ values are reported in nM molecules for all assays.

*M13 mp18 ssDNA* was prepared as described (49). The concentration of DNA was determined spectrophotometrically using an extinction coefficient of 8, 780 M^−1^cm^−1^ (nucleotides). Purified DNA was stored in small aliquots at −80°C.

ϕX174 ssDNA was purchased from New England Biolabs. The concentration of DNA was determined spectrophotometrically using an extinction coefficient of 8, 780 M^−1^cm^−1^ (nucleotides). Following concentration determination, the ssDNA was distributed into small aliquots and stored at −80°C.

*Model fork-DNA substrates* consisting of a homologous core of 12 bp flanked by heterologous duplex arms of 19-25 bp were constructed by annealing gel-purified oligonucleotides. The substrate design was similar to that used previously for 3 and 4-strand substrates (49,52)}. The junction point can branch migrate within the homologous core, whereas the heterologous arms prevent the spontaneous resolution of the junction DNA.

Model fork substrates were prepared by annealing six oligonucleotides in various combinations: PB170 (5’-CTAGAGACGCTG CCGAATTCTGGCTTGGATCTGATGCTGTCTAGAGGCCTCCACTATGAAATCGCTGCA-3’), PB171 (5’-GCGATTTCATAGTGGAGGCCTCT AGACAGCA-3’), PB172 (5’-TGCTGTCTAG AGACTATCGATCTATGAGCTCTGCAGC-3’), PB173 (5’– CCGGGCTGCAGAGCTCATAGA TCGATAGTCTCTAGACAGCATCAGATCCAAGCCAGAATTCGGCAGCGTCT-3’), PB345 (5’-GCGATTTCATAGTGGAGGCCTCTAGACAGCACGCCGTTGAATGGGCGGATGCTAATTACTATCTC) and PB346 5’-GAGATAGTAATT AGCATCCGCCCATTCAACGGCGTGCTGTCTAGAGACTATCGATCTATGAGCTCTGCAGC). Each oligonucleotide was purified using denaturing polyacrylamide gels containing 6 M urea. Purified oligonucleotides (1 μM each in molecule) were annealed in a total volume of 50 μl containing incubation at 100°C for 5 min, followed by cooling to room temperature overnight. The extent of annealing was verified by non-denaturing PAGE using 5’-end labeled oligonucleotides annealed under identical conditions (data not shown). Junctions were added directly to ATPase assays without further purification. As the annealing reactions contained 10 mM magnesium acetate, the concentration of magnesium ions in each assay was adjusted accordingly.

### Proteins

*RecG* protein was purified as described previously (52). The protein concentration was determined spectrophotometrically using an extinction coefficient of 49,500 M^−1^ cm^−1^ (71).

*His-PriA* cloning was described previously (38). To lyse cells, a 1L culture was grown with protein expression induced by the addition of 500μM IPTG at an OD_600_ of 0.5, followed by growth for an additional 3 hours. Cells were harvested by centrifugation and lysis of the resuspended cell pellet was initiated by the addition of lysozyme (1 mg/ml final), and benzonase (3 μl), followed by stirring for 30 minutes at 4°C. Deoxycholate was added to 0.05% final, and the mixture stirred for an additional 30 minutes. Imidazole and KCl were added to a final concentration of 30 and 600 mM, respectively. The whole-cell lysate was centrifuged at 37,000×G at 4°C for 1 hour. The cleared cell lysate was loaded onto a 5 ml HisTrap FF column equilibrated in Binding Buffer (30 mM Imidazole; 15.4 mM Na_2_HPO_4_; 4.5 mM NaH_2_PO_4_; 600 mM KCl; (pH 7.4)). The nickel column was subjected to three washes sequentially: Binding Buffer (50 column volume (CV)), Binding Buffer with 0.2% NP40 (40 CV), Binding Buffer (30 CV). Proteins were eluted using a linear, imidazole gradient (30 mM to 500 mM in the same buffer). Fractions containing PriA were identified by 12% SDS-PAGE, pooled, and dialyzed overnight into heparin column binding buffer (20 mM Tris-acetate, (pH 7.5), 0.1mM EDTA, 1mM DTT, 40mM KCl, and 10% glycerol).

The next day, the dialyzed protein was subjected to centrifugation at 10,000xG for 10 minutes and the supernatant was applied to a 20 ml heparin FF column equilibrated in heparin column binding buffer. Following a wash to baseline, the protein was eluted with a linear gradient (10 column volumes) from 40-1,000mM KCl in the same buffer. Fractions containing PriA (and no detectable contaminants) were identified by SDS-PAGE, pooled and dialyzed overnight against storage buffer (20 mM Tris-HCl pH 7.5; 0.1mM EDTA, 1 mM DTT, 100 mM NaCl, 50% glycerol). Protein concentration was determined spectrophotometrically using an extinction coefficient of 104,850 M^−1^ cm^−1^ (71). The presence of the N-terminal histidine tag did not alter the activities of the protein relative to the untagged version (data not shown).

*SSB proteins –* the single-stranded DNA-binding protein (SSB) was purified from strain K12ΔH1Δtrp as described (72). The concentration of purified SSB protein was determined at 280 nm using ε = 30, 000 M^−1^ cm^−1^. The site size of SSB protein was determined to be 10 nucleotides per monomer by monitoring the quenching of the intrinsic fluorescence of SSB that occurs on binding to ssDNA, as described (73). His-SSB was purified as described previously (38). His-SSBΔC8 was purified as described (66).

*Bacteriophage gene 32 protein (gp32)* was over-expressed and purified as described (74,75). The concentration of purified gp32 was determined at 280 nm using ε = 37,000 M^−1^ cm^−1^ (76). The site size of gp32 was determined to be 7 nucleotides per monomer by monitoring the quenching of the intrinsic fluorescence of gp32 that occurs on binding to ssDNA, as described (76).

### ATP hydrolysis assay

The hydrolysis of ATP was monitored using a coupled spectrophotometric assay (49,52). The standard reaction buffer contained 20 mM Tris-acetate (pH 7.5), 1 mM DTT, 0.3 mM NADH, 7.5 mM PEP, 20 U/mL pyruvate kinase, 20 U/mL lactate dehydrogenase, 10 nM PriA, 1 mM ATP, and 10 mM magnesium acetate (but varied according to the DNA cofactor present). The rate of ATP hydrolysis was calculated by multiplying the slope of a tangent drawn to linear portions of time courses by 159. In a typical reaction, close to 200 data points were used to draw a linear fit to the data to calculate reaction rates. In assays with SSB present, it was stoichiometric relative to the fork. For example, for 100nM fork 1, 200nM tetramer was required (one per ssDNA arm); for forks 2 and 3, only 100nM tetramer was required.

To obtain kinetic parameters, data were analyzed using non-linear curve fitting in Prism v 8.4.3 (GraphPad Software, LLC). DNA titration data were fit to the Hill equation (V = (V_max_.[DNA]^n^) / ([S_0.5_]^n^ + [DNA]^n^) or the Michaelis-Menten equation (V = (V_max_.[DNA])/(K_m_+ [DNA]) (77). ATP titration data were fit to the Michaelis-Menten equation only. In situations where the binding appeared cooperative, a comparison was done in Prism to determine which model more accurately described the data. Here models were compared using the Comparison of fit function and models discriminated using both F-test and P-values. In those instances where the Hill equation more accurately described the data, P values <0.0001 and high F-values were obtained (data not shown).

In salt-titration experiments, the same reaction buffers were used (see above). Reactions were initiated by the addition of either PriA or RecG following a 5-minute incubation of all other components. When single-strand binding proteins were present, they were added before PriA or RecG at the concentrations indicated in figure legends. Once a steady-state rate of ATP hydrolysis was achieved, NaCl was added in 12.5 mM increments (1μl volumes). This was repeated until all ATP hydrolysis of either PriA or RecG ceased. The resulting hydrolysis rate in each steady-state region was calculated and expressed as a percent of the steady-state rate in the absence of NaCl. The total volume used to calculate the final concentration of NaCl was adjusted after each addition to correct for the additions themselves. A line of best fit was drawn for data points between each addition, to obtain the ATP hydrolysis rate after each salt increment. The average number of data points used to determine the reaction rate was 14. These rates were subsequently graphed to determine the concentration of NaCl resulting in a 50% reduction in the rate of ATP hydrolysis which corresponds to the salt-titration mid-point.

## Author contributions

Tan did the experiments. Tan and Bianco analyzed the data. Bianco wrote the manuscript

## Funding and additional information

Funding in the Bianco laboratory is supported by NIH grant GM10056 to PRB. “The content is solely the responsibility of the authors and does not necessarily represent the official views of the National Institutes of Health.”

## Conflict of interest

The authors declare there are no conflicts of interest.

---

**Supplementary Figure 1.**
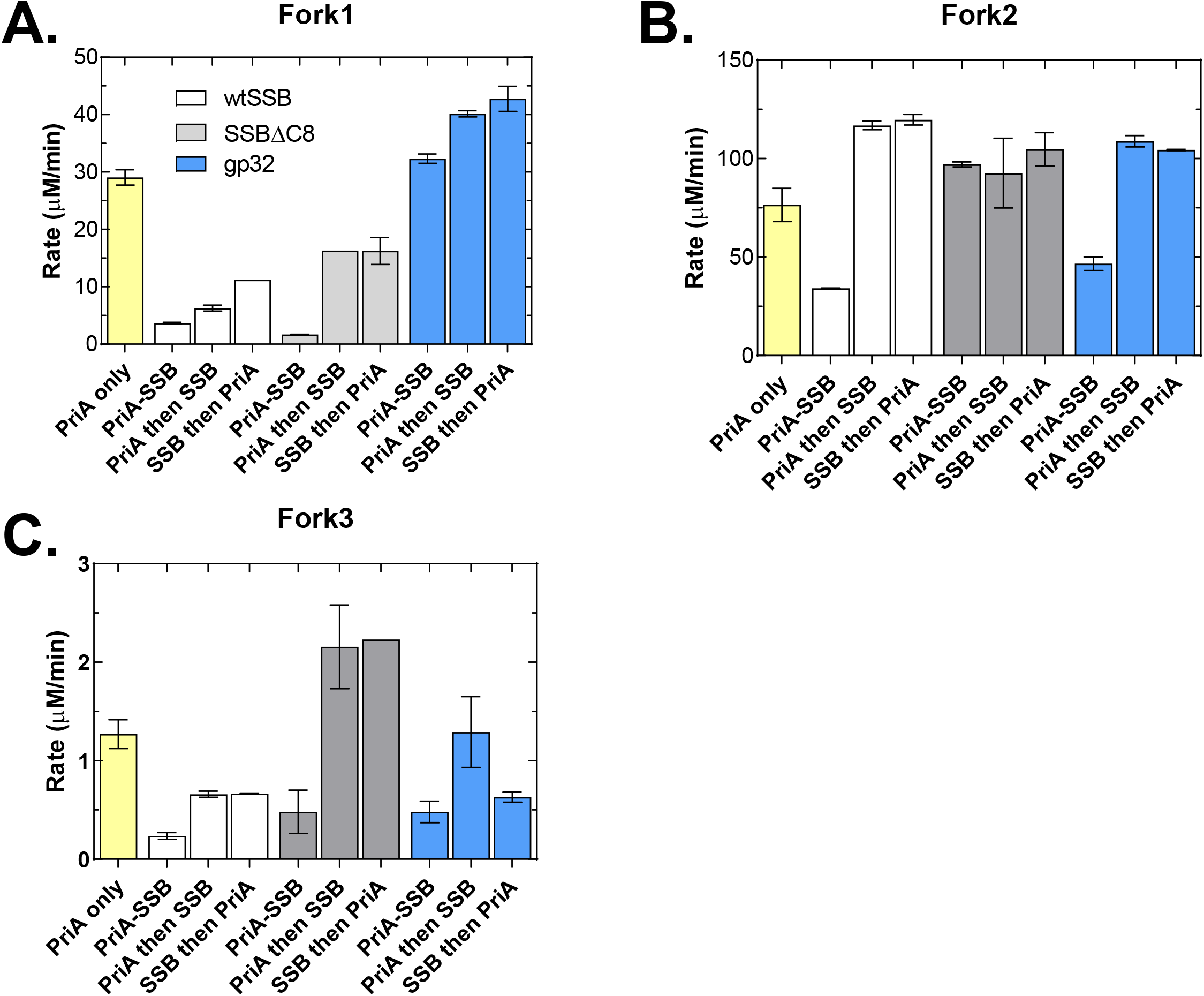
The order of addition at forks is critical to the stimulation of PriA afforded by SSB. Assays were done as descried in the Experimental Procedures. Assays contained 100nM fork DNA, 10nM PriA and ssb proteins indicated. For SSB and SSBΔC8, they were present at 200nM tetramer for fork 1 and 100nmM tetramer for forks 2 and 3. Gp32 was present at the same concentrations but in monomer. For ssb-PriA assays, proteins were mixed on ice for 30 minutes prior to being added to assays. A minimum of 2 assays per protein condition per fork were done. For some data sets, the data were so close it appears as if no errors bars are present.

**Supplementary Figure 2.**
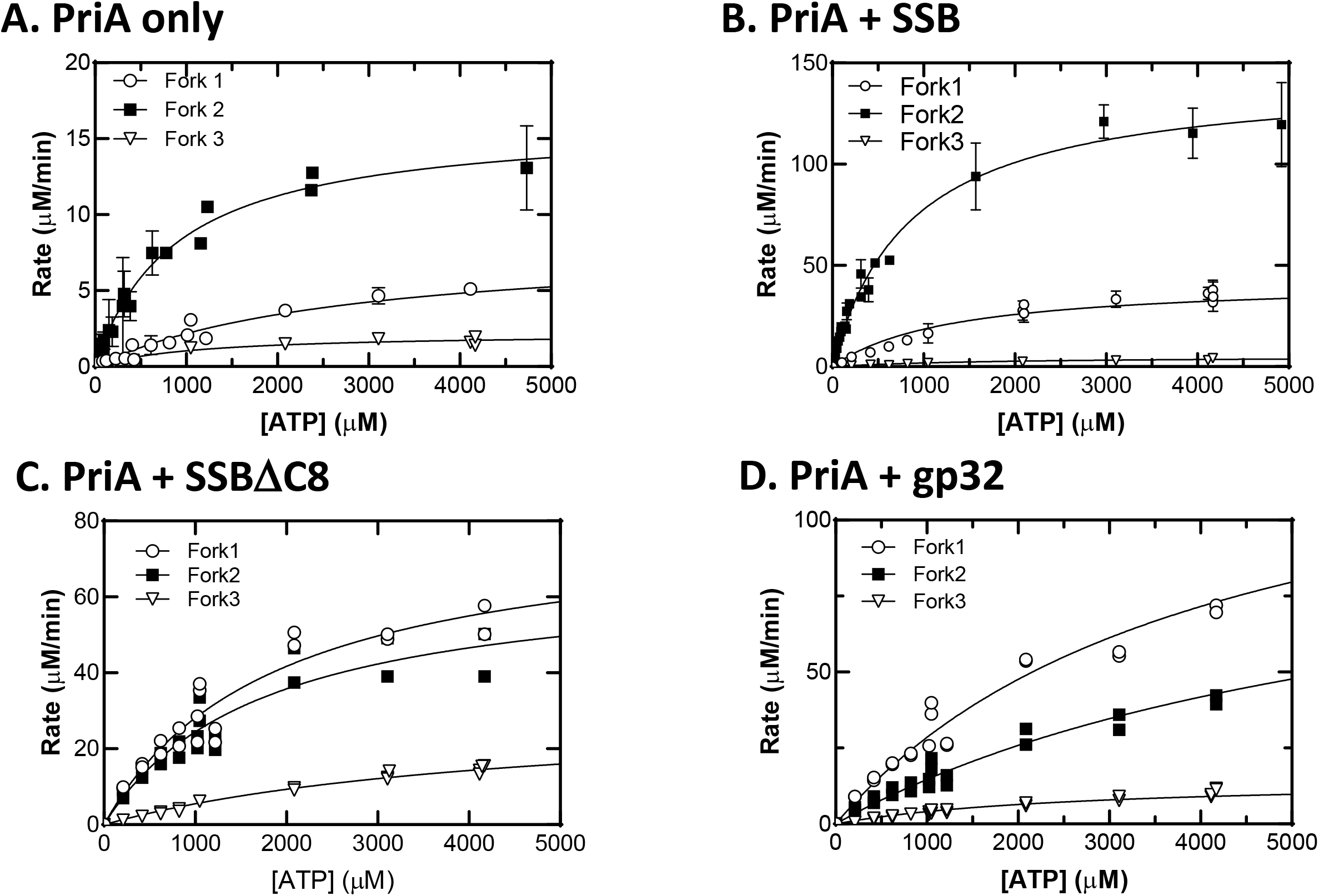
Single strand binding proteins have DNA substrate-dependent effects on the ATPase activity of PriA. Assays were done as descried in the Experimental Procedures. Assays contained 10mM MgOAc, 100nM fork DNA, 10nM PriA and ssb proteins indicated. For SSB and SSBΔC8, they were present at 200nM tetramer for fork 1 and 100nmM tetramer for forks 2 and 3. Gp32 was present at the same concentrations but in monomer. For ssb-PriA assays, proteins were mixed on ice for 30 minutes prior to being added to assays. Two – five assays per [ATP] per fork were done on separate days. Data were fit to the Michaelis-Menten Equation.

**Supplementary Figure 3.**
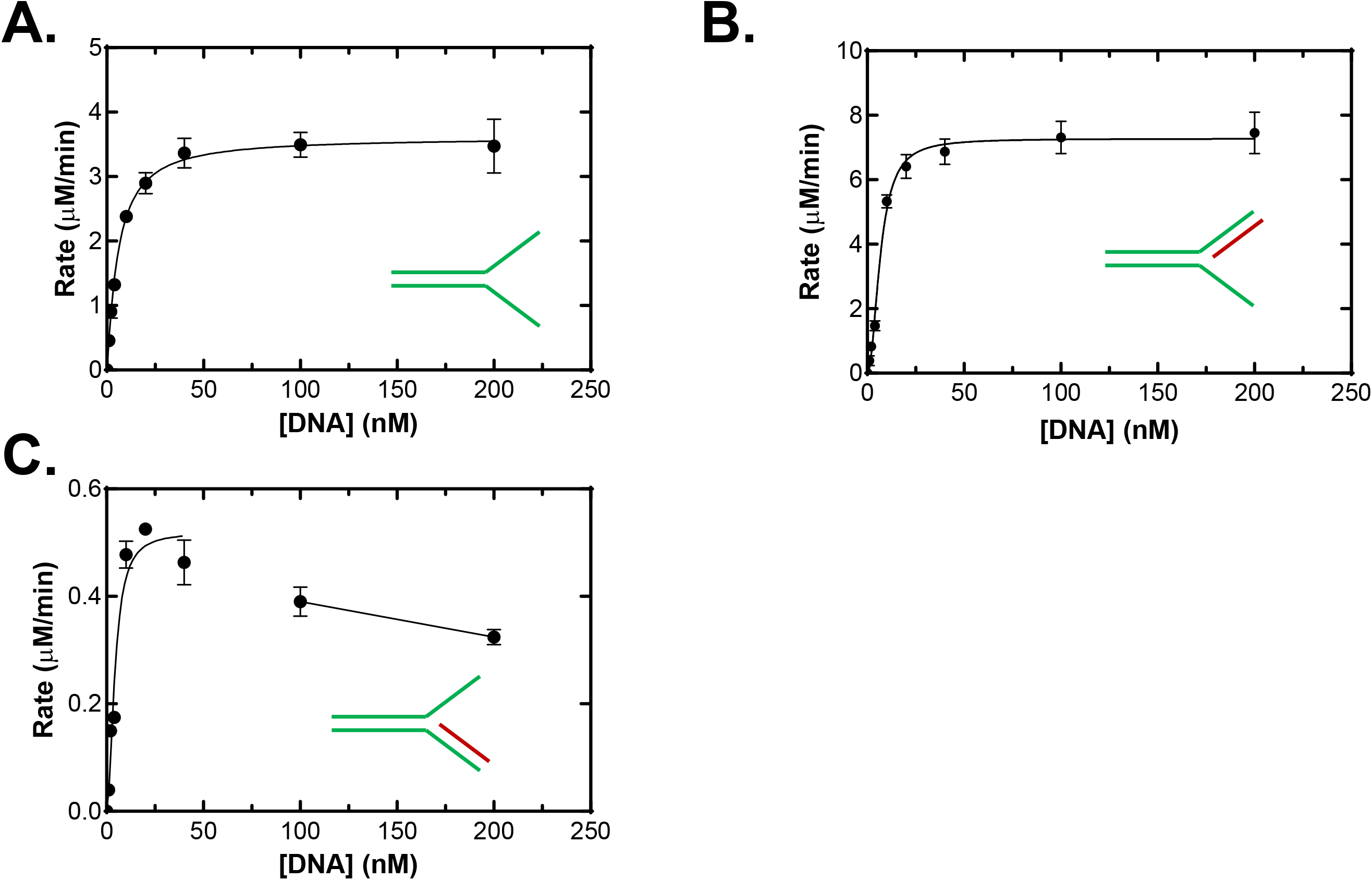
DNA titrations reveal surprises into PriA DNA cofactor preference. Assays were done as descried in the Experimental Procedures. Assays contained 10mM MgOAc, 1mM ATP, 10nM PriA and forks as indicated. Two – five assays per [DNA] were done on separate days. Data were fit to the Hill Equation. The concentration of DNA is reported in nM molecules. (A), fork 1; (b), fork 2 and (C), fork 3. For the analysis of fork3, only data at or below 50nM DNA were used in calculations.

